# From smartphone to gene: Automated image-based phenotyping captures flowering diversity of common buckwheat

**DOI:** 10.64898/2026.11.03.763633

**Authors:** Carles Quesada-Traver, Fabian Hess, Flavian Tschurr, Lukas Kronenberg, Gabriele Rimat, Steven Yates, Bruno Studer

**Affiliations:** Molecular Plant Breeding, Institute of Agricultural Sciences, ETH Zurich, Universitaetstrasse 2, 8092 Zurich, Switzerland; Grassland Sciences, Institute of Agricultural Sciences, ETH Zurich, Universitaetstrasse 2, 8092 Zurich, Switzerland; Precision Phenotyping, Institute of Agricultural and Nutritional Science, Martin Luther University Halle-Wittenberg, Theodor-Lieser-Strasse 11, 06120 Halle (Saale), Germany; Institute for Crop Science and Plant Breeding, Bavarian State Research Center for Agriculture, LfL, Am Gereuth 2, 85354 Freising, Germany; Puregene AG, Etzmatt 273, 4314 Zeiningen, Switzerland

## Abstract

Common buckwheat (*Fagopyrum esculentum* Moench) exhibits an indeterminate flowering habit, resulting in the simultaneous occurrence of flowering, seed development and seed ripening. This contributes to yield losses and complicates mechanized harvesting. Therefore, the development of varieties with a desirable flowering pattern is a breeding priority in buckwheat. However, phenotyping of buckwheat flowering under field conditions remains challenging because manual assessments are time consuming and expertise dependent. Here, we show that imaging devices such as smartphones or cameras can be used for phenotyping of flowering traits in a buckwheat collection encompassing 169 accessions. Flowering traits describing flowering time (*n*=10), flowering intensity (*n*=6) and abscission (*n*=6) were extracted from 37,078 images captured over five growing seasons using a Mask R-CNN segmentation model combined with statistical modelling of flowering dynamics. The 22 extracted traits were highly heritable (0.5 < ℎ^2^ < 0.93) and correlated well (0.84 < *r* < 0.94) with manually assessed flowering stages while providing additional information on flowering intensity. Genome-wide association studies revealed 171 genomic regions associated with flowering traits, including marker-trait associations for agronomically important breeding targets. Candidate genes had functional annotations including “Auxin response factor” and “Ethylene-insensitive protein”, gene families involved in hormone signalling and developmental regulation, controlling flowering in other species. These results demonstrate how digital imaging combined with automated feature extraction can be leveraged to gain insight on buckwheat flowering patterns in large scale field trials across different phenotyping platforms. This paves the way to breed for desired flowering patterns using molecular tools in a targeted manner.

## 1. Introduction

Common buckwheat (*Fagopyrum esculentum* Moench), hereafter referred to as buckwheat, is a neglected crop having the potential to diversify cereal-dominated farming systems (Joshi et al., 2019). Its resistance to most pests and diseases (Campbell, 1997), its short growing season and the possibility to be managed using machinery make buckwheat a versatile and valuable addition to most crop rotations (Farooq et al., 2016; Strahm et al., 2019). In addition, buckwheat offers a source of gluten-free starch and a favourable protein composition for human consumption (Škrabanja & Kreft, 2016). However, one major issue that limits buckwheat adoption is its prolonged flowering phase (Campbell, 1997).

This flowering behaviour complicates mechanical harvesting of buckwheat, as prolonged and asynchronous flowering results in the simultaneous presence of flowers and seeds over an extended period. This disrupts harvest operations and reduces the yield potential (Campbell, 1997; Joshi et al., 2019). Early harvest results in the presence of flowers, unripe seeds and high moisture contents in the produce. Late harvest increases the risk of seed scattering and pre-harvest sprouting, resulting in yield losses (Funatsuki et al., 2000; Morishita et al., 2020; Ohsawa, 2020). In contrast, prolonged flowering is beneficial from an ecological perspective, as it provides nectar and pollen for insects (Cawoy et al., 2009; Strahm et al., 2019). Thus, the lack of buckwheat accessions with flowering patterns tailored to their role in the cropping system hinders increased farmer adoption and the exploitation of its ecological benefits (Hess et al., 2026b).

The use of genomic methods to breed for specific flowering traits could accelerate buckwheat improvement, as suggested for other neglected crops (Varshney et al., 2012; Joshi et al., 2019; Yabe and Iwata, 2020; Nature Communications Editorial Staff, 2024). However, using such methods requires efficient phenotyping of buckwheat flowering dynamics. Although BBCH scales have been developed to describe flowering patterns in buckwheat (Arduini et al., 2016; Grimes et al., 2025), their application is laborious and difficult to perform in accessions with high diversity. Stage assignment is not straightforward and requires experience, as developmental stages overlap and vary within plots. As a result, BBCH-based phenotyping does not scale well to large trials. Moreover, BBCH scales do not capture traits related to flowering intensity (i.e. number of flowers or flowering extent). Therefore, efficient and high-throughput approaches are needed to overcome this phenotyping bottleneck (Cobb et al., 2013).

In recent years, instance segmentation with deep learning has become a common approach to automate crop phenotyping by detecting and quantifying plant structures directly from images (Murphy et al., 2024). Mask R-CNN, in particular, has been widely adopted because it provides object-level masks that support counting and area-based measurements in complex scenes (He et al., 2017). For example, Mask R-CNN has been used to segment on-tree fruits to enable automated monitoring and size estimation in orchards (Giménez-Gallego et al., 2024). At the other end of the scale, fine-tuned Mask R-CNN models have also been applied to microscopic images for the estimation of stomatal density in apple trees (Zuffa et al., 2025). Image-based modelling approaches have also been used to characterize flowering behaviour. For example, Oppliger et al. (2025) used image-based flower detection to quantify flowering dynamics in peas (*Pisum sativum* L.), revealing differences between breeding lines and correlations between flower number and yield. In wheat (*Triticum aestivum* L.), Wang et al. (2019) used an image-based classifier to measure the variation in heading date across breeding lines and linked this variation to genomic regions controlling the trait. These examples show that modelling dynamic flowering patterns from image-based features can quantify phenology and link trait variation to genomic regions.

In this study, we developed an image-based, automated pipeline to phenotype buckwheat flowering traits relevant for breeding. To ensure that our method can be adapted to different contexts and the equipment available in multi-environment trials we compare the performance of different imaging devices, including smartphones, in different environments. Genome wide association studies are performed on the traits extracted from the image-based pipeline and manual grading of BBCH stage to compare and cross validate the results. In addition, the identified marker-trait associations provide a genetic basis for the targeted breeding of flowering dynamics in buckwheat breeding.

## 2. Materials and methods

### 2.1. Experimental design, image acquisition and germplasm collection

Field trials were conducted in 2023 and 2024 to test the 169 buckwheat accessions of the ETH Zurich/BLW buckwheat collection described in Hess et al. (2026c) at the Plant Research Station of ETH Zurich in Eschikon, Switzerland (47.449 N, 8.682 E, 520 m a.s.l.) and at the Experimental station Baumannshof of the Bavarian State Research Center for Agriculture (LfL) (48.701 N, 8.531 E, 360 m a.s.l.). A total of five field trials were conducted including four trials with buckwheat as the main crop (ETH23, ETH24, LfL23, LfL24) and one late-sown trial with buckwheat as the second crop which was located at ETH (ETH24L) (Figure 1A). All field trials followed an augmented row-column design, with replicated accessions in complete blocks and spatially distributed check accessions to control within-field heterogeneity. The ETH trials (ETH23 and ETH24) comprised 169 accessions, with four of these accessions serving as checks. The LfL trials also comprised 169 accessions with seven and one of these accessions serving as checks in 2023 and 2024, respectively. Accessions were replicated three times in both ETH trials, while one and two replications were included in the LfL trials in 2023 and 2024, respectively. A subset of the collection comprising 55 accession was grown in ETH24L in two replicates, with one accession serving as spatial check(Table 1). The image acquisition in Eschikon was performed using a prototype of LITERAL® (Hyphen, Avignon, France). At Baumannshof, images were taken using a Samsung Galaxy S10 smartphone (Samsung Electronics, Suwon, South Korea) in 2023 and an Olympus digital camera model TG-6 (Olympus Corporation, Tokyo, Japan) in 2024. Both devices were attached to a bamboo stick (1.5 m) and the camera height was adjusted according to the maximum canopy height in the plots at the beginning of each imaging campaign. All imaging devices were used with auto settings (focus, ISO, aperture and shutter speed). During dry weather conditions, images were taken twice per week throughout the growing season from germination to harvest. Image acquisition was suspended during rain if it interfered with equipment handling or image quality. Across experiments, plots were imaged between 7 and 35 times (median 9 to 31), spanning 65 to 121 days between the first and last image (SD 5.5 to 21.8 days; Table 1).

**Figure 1.**
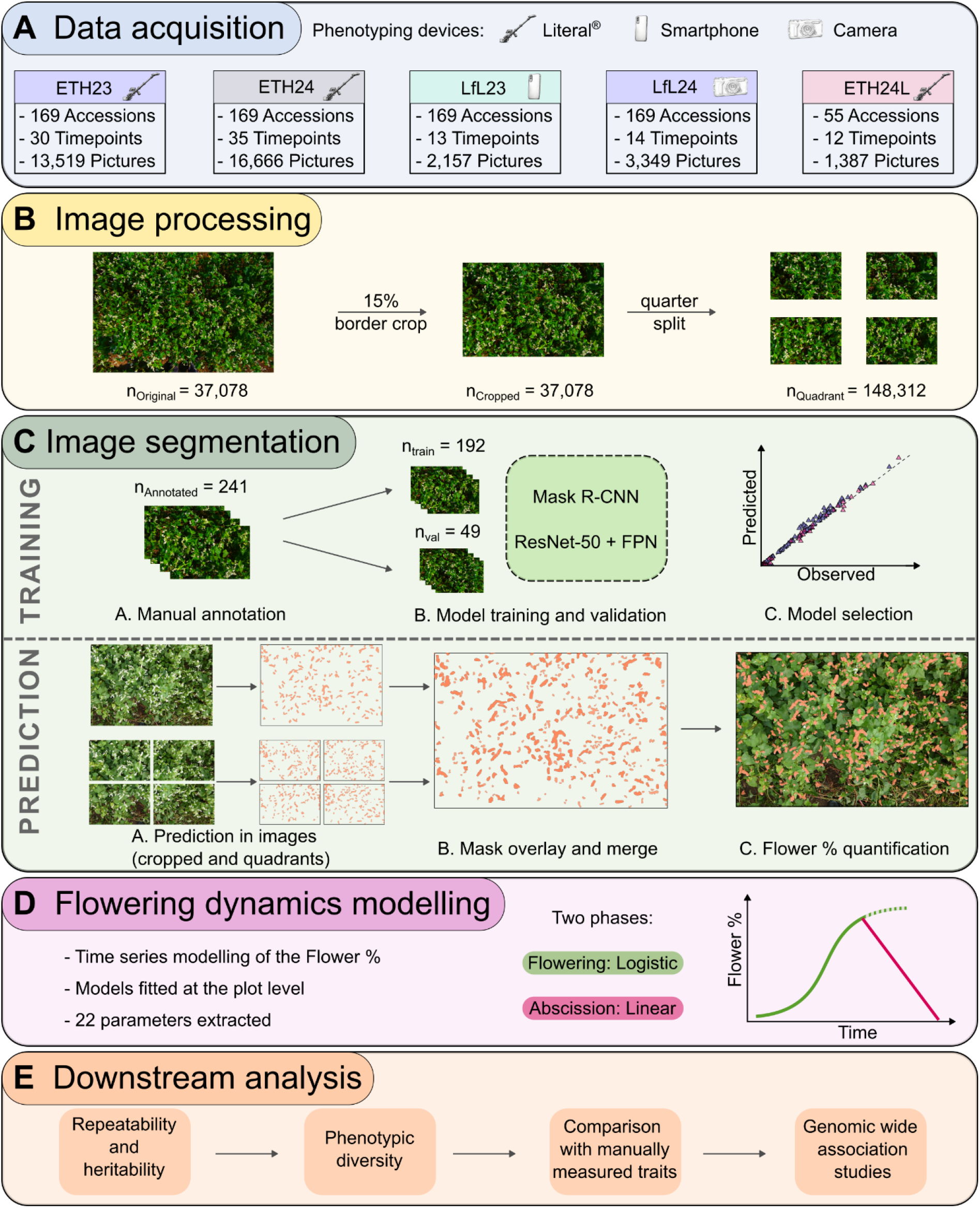
Overview of the image-based flower detection phenotyping method and downstream analysis. A. depicts details on data acquisition such as imaging device (i.e. Literal^®^, smartphone and camera), number of accessions, timepoints and images taken in each of the trials. B. illustrates the image preprocessing pipeline which includes border cropping and image splitting in quadrants. C. shows the image segmentation modelling strategy. For training, the number of quadrants in the training (*n_train_*) and validation (*n_val_*) splits are shown. For prediction, the strategy of predicting in the cropped image as well as in the quadrants with the subsequent mask overlaying and merging is illustrated for one instance. In D. the two-phase flowering dynamics modelling is shown, where the three-parameter logistic function describing flowering and the linear function describing abscission are shown in green and magenta, respectively. The *x* axis represents the timepoints and the *y* axis the percentage of pixels in the image assigned as flower (Flower %). In E. the steps of downstream analysis are illustrated.

**Table 1.**
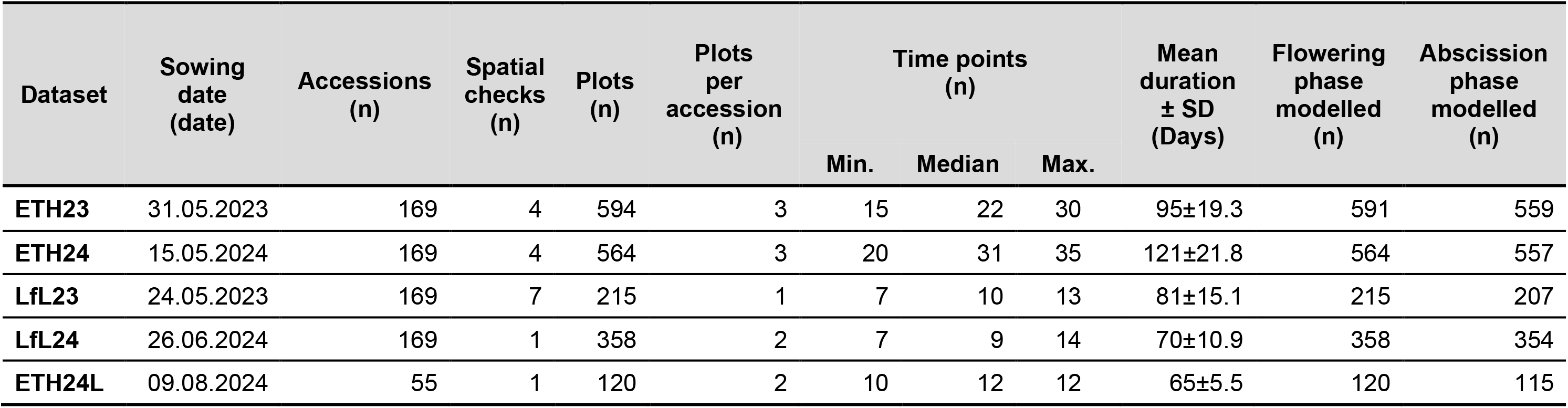
Description of the field experiments and data used for flowering dynamics modelling. For each experiment, the sowing date, the number of accessions (including spatial checks)and plots planted in the field as well as the median number of plots per accession (excluding checks) are reported. Additionally, the maximum, median and minimum number of time points at which images were taken per plot are depicted. The mean duration and standard deviation between the first and the last image taken is also shown. Finally, thenumber of plots successfully modelled in terms of flowering andabscission phase are also provided.

The genetic characterization of the 169 buckwheat accessions based on genome-wide allele frequency fingerprints, as well as the marker filtering procedure applied, were described in Hess et al. (2026c). In short, one hundred leaves of individual plants per accession were sampled in 2023 and 2024 to form two pooled samples for each accession. From these samples, DNA was extracted to construct double digest restriction-site associated DNA libraries for sequencing. The sequencing data from both years was pooled (resulting in a single dataset per accession) for subsequent calling of single nucleotide polymorphisms (SNPs) and calculation of allele frequencies representative of the genetic composition of each accession. This resulted in a matrix of 52,182 SNP frequencies after filtering for a minor allele frequency greater than 0.05. Missing values were imputed through linkage-informed k-nearest neighbour-derived weighted mean allele frequency prediction using imputef (v.1.1.1; Paril et al., 2025).

### 2.2. Phenology

The plant development stage of each plot for the ETH23 and ETH24 trials was assessed twice a week during the growing season. A selection of development stages as described in a proposed BBCH scale for buckwheat was used (Arduini et al., 2016). Specifically, stages 00 (Seed), 10 (Cotyledon), 11 (Leaf 1), 12 (Leaf 2), 13 (Leaf 3), 14 (Leaf 4), 50 (Blossoming), 60 (First Flower Open), 62 (Flower Open Terminal Inflorescence, TINF), 70 (Green Fruits), 85 (Brown Fruits), 67 (All Fruits Brown TINF), 86 (End Flowering TINF) of the proposed BBCH scale were assessed. The development stages were numbered (1-13) in the order specified before to account for the temporal sequence of buckwheat development. A monotonically increasing *p*-spline was fitted to the data of each plot using the *scam* function of the R package scam (v.1.2.17) with parameters k = 8 and bs = “mpi” to interpolate between recorded stages. Daily BBCH stages were predicted from the splines and used for further analysis. Data of stages 60, 62, 70 and 85 have been published by the authors in a previous preprint manuscript but are presented here again for comparison with image-derived flowering traits (Hess et al., 2026b).

### 2.3. Modelling and data analysis

Plot-level flowering traits were derived by segmenting flowers in each image and subsequent modelling over time to quantify flowering dynamics for downstream analyses (Figure 1). All calculations were performed in R (v.4.4.0) unless stated otherwise.

#### 2.3.1. Image processing and flower segmentation

Each original image was pre-processed by applying a fixed border crop of 15% to exclude non-plot regions from the picture. The cropped images were then divided into four quadrants (see Figure 1B). A set of 241 quadrants from randomly chosen images taken in the ETH23 field trial spanning the growing season was selected, and flower clusters were manually annotated using the polygon tool on the VGG Image Annotator (VIA) v.3.0.11 (Dutta & Zisserman, 2019). This set was randomly divided into two groups, which constituted the training (*n* = 192) and the validation (*n* = 49) sets, respectively. Mask R-CNN with a ResNet-50 backbone and a Feature Pyramid Network (FPN) initialized with pre-trained COCO weights (Lin et al., 2014) was used for flower instance segmentation. The model was trained for a total of 1,200 epochs using stochastic gradient descent with momentum (µ = 0.9). The loss function to be minimized, combined classification and bounding box regression losses for both the Region Proposal Network (RPN) and the Fast R-CNN head, along with a mask prediction loss for the Mask R-CNN head, regularized by L2 weight decay (*λ* = 1 × 10*^−^*^4^), as described in Faster R-CNN (Ren et al., 2017) and Mask R-CNN (He et al., 2017).

The base learning rate was set to 0.01 with a short linear warm-up and subsequent step decays. An effective batch size of approximately eight was achieved through gradient accumulation. In addition, model checkpoints were saved every 50 epochs and used to calculate metrics on the validation set. Training was performed using TensorFlow (v.1.15.5) and TensorPack (v.0.11.0) in Python (v.3.8.13) running on a single NVIDIA GeForce RTX 3080 Ti (NVIDIA Corporation, Santa Clara, CA, USA). The best checkpoint, based on training and validation metrics, was selected and retained for further predictions.

Once selected, the model was used to generate the prediction masks of the flower clusters in all the collected images. For each image, predictions were produced on the cropped image and on each of the four quadrants to improve detection across different scales. Quadrant outputs were mapped back onto the cropped image, and the predictions were combined into a single collection of polygon masks per image. Any overlapping or duplicated masks were merged into single masks to yield one final set of masks per image. Finally, for each image the percentage of pixels inside the masks relative to the total number of pixels was calculated and used as a proxy for flowering intensity in subsequent analyses and is hereafter referred to as Flower %. Image and mask manipulation were performed using functions from the Python packages cv2 (v.4.6.0), shapely (v.9.2.0) and PIL (v.2.9.6).

#### 2.3.2. Flowering dynamics modelling and parameter extraction

To extract flowering-related phenotypes at the plot level, the observations of each time series were imputed to daily resolution using cubic smoothing splines, fitted with the smooth.spline function from the stats package (v.4.5.1) in R. The measurement period was then divided into two phases: flowering and abscission. The flowering phase was defined by the start of the measurements and the time point of maximum flower coverage, while the abscission phase extended from this point to the end of the measurement period. A diagram showing how this sequential model is fitted, along with all the extracted parameters, is illustrated in Figure 2 and a recollection of all the traits, with information on their calculation and type can be seen in Table S1. All date-related parameters were normalized to days after sowing (DAS). The flowering phase was modelled using a three-parameter self-starting logistic function described in Equation 1:

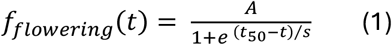

**Figure 2.**
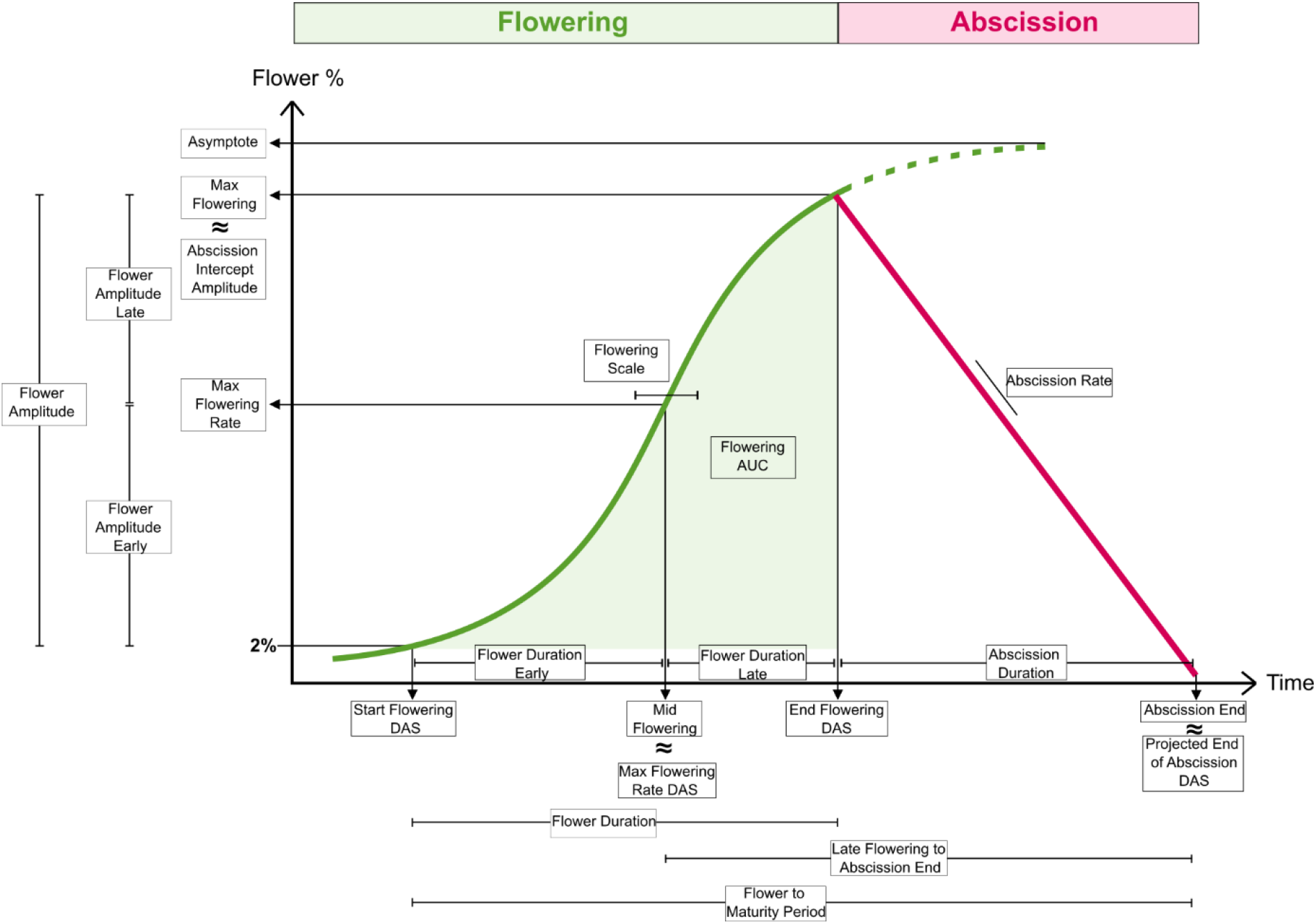
Representation of the two-phase modelling strategy showing the three-parameter logistic function (green) modelling the flowering phase and the linear model (magenta) modelling the abscission phase. Additionally, all the parameters extracted and calculated from the fits are displayed in boxes.

where *f_flowering_*(*t*) represents the Flower % at time *t* (days), *A* is the asymptote maximum flower coverage (%), *t*_50_ denotes the inflection point corresponding to mid flowering (days) and *s* is a scaling parameter controlling the steepness of the curve (days). The model was fitted by nonlinear least squares using the nls function with the SSlogis formulation from the R package stats (v.4.5.1). From each fitted curve, the coefficients *A*, *t*_50_ and *s* were retrieved as Asymptote (%), Mid Flowering (days) and Flowering Scale (days). Based on the sensitivity of the image segmentation model, Start Flowering DAS (days) was defined as the first date when the percentage of flowers covered at least 2% of the image area. The first derivative of the fitted curve, *df_flowering_*(*t*)/*dt*, was used to identify the time point at which the instantaneous rate of increase in flower coverage was maximal, this time point was defined as Max Flowering Rate DAS (days), and the flower coverage at this time point was recorded as Max Flowering Rate (%). Following an approach similar to that used to characterize senescence dynamics in wheat (Tschurr et al. 2024), the second derivative of the fitted logistic function, *d*^2^*f_flowering_*(*t*)/*dt*^2^, was used to determine End Flowering DAS (days) and Max Flowering (%), corresponding to the point at which the acceleration of flowering declines towards the stationary phase of the curve. Finally, the area under the fitted logistic curve (Flowering AUC, % days) was calculated by numerical integration of the fitted logistic function over the flowering phase. The abscission phase was modelled using a linear function of time as described in Equation 2:

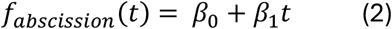

where *f_abscission_*(*t*) represents the Flower % at time *t* (days). The model was fitted with ordinary least squares using the lm function from base R. The estimated coefficient *β*_1_ was interpreted as Abscission Rate (% days^-1^), while *β*_0_ represented the Abscission Intercept Amplitude (%). The point at which the fitted line crossed the time axis (*f_abscission_*(*t*) = 0) during the abscission phase was recorded as Abscission End (days), from which a normalized value (Abscission Projected End DAS (days)) was also calculated.

In addition to the parameters directly obtained from the flowering and abscission models, six duration parameters and three amplitude parameters were calculated. Flower Duration (days) was described as the interval from Start Flowering DAS to End Flowering DAS. This duration was further divided into Flower Duration Early (days), spanning from Start Flowering DAS to Max Flowering Rate DAS, and Flower Duration Late (days), from Max Flowering Rate DAS until End Flowering DAS. Abscission Duration (days) was defined as the interval between End Flowering DAS and Abscission Projected End DAS. Extending this timeframe, the Flower to Maturity Period (days) was defined as the interval between Start Flowering DAS and Abscission Projected End DAS. While Late Flowering to Abscission End (days) was defined as the period between Max Flowering Rate DAS and Abscission Projected End DAS. Changes in flower coverage during the flowering phase were quantified using three amplitude parameters. Flower Amplitude (%) was calculated as the difference between Max Flowering and Start Flowering (2%). This total change was further divided into Flower Amplitude Early (%), defined as the difference in flower coverage between Max Flowering Rate and Start Flowering (2%), and Flower Amplitude Late (%), defined as the difference between Max Flowering and Max Flowering Rate.

To assess the goodness-of-fit, the *R*^2^ statistic was calculated from the sum of the squared errors for the fit of the flowering and abscission phases separately. Parameters extracted from these fits with a *R*^2^ lower than 0.5 were excluded from the subsequent analysis.

#### 2.3.3. Spatial correction and repeatability

The traits extracted at the plot level by the flowering dynamics model were further analysed using a two-stage linear mixed model as described in (Roth et al., 2021) and (Roth et al., 2024). In the first stage, adjusted genotype means of all traits were calculated for each year-site using the R package SpATS (v.1.0.19) (Rodríguez-Álvarez et al., 2018). The model parameters were specified as:

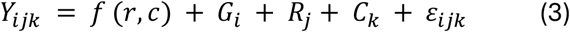

where *Y_ijk_* is the parameter value of the plot in the *i^t^*^ℎ^ genotype, *j^t^*^ℎ^ year-site and of the *k^t^*^ℎ^replication, *f* (*r*, *c*) is a smooth bivariate surface defined over rows and ranges, *G_i_* is the effect of the *i^t^*^ℎ^ genotype while *R_j_* and *C_k_* are the effects of the *j^t^*^ℎ^ row and *k^t^*^ℎ^ range, respectively. The plot residuals are denoted as *ε_ijk_*. In Equation 3, *G_i_* was set as random to calculate repeatability (Oakey et al., 2006), while for the calculation of best linear unbiased estimators (BLUEs) the term was set as fixed. Moreover, the diagonal of the inverse of the variance-covariance matrix from this model was extracted and used as weights in the next stage.

#### 2.3.4. Genotype means across year-sites and heritability

To calculate genotype means across year-sites, the R package ASReml (v.4.1.0.176) was used to specify the model:

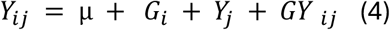

where *Y_ij_* are the BLUEs calculated in the previous stage, µ the global intersect, *G_i_* the genotype effect, *Y_j_* the year-site effect and *GY _ij_* represents the weighted genotype by year-site interaction modelled through a diagonal error variance structure for genotypes nested within year-sites. Since we used adjusted genotype means per year-site, the GxE interaction is confounded with the residual error and no explicit quantification of this variance component can be given. The *G_i_* term in Equation 4 was set as random to calculate BLUPs and heritabilities (Cullis et al., 2006) while it was specified as fixed for the calculation of BLUEs across years.

#### 2.3.5. Correlations between phenotypic values

Pairwise Pearson correlation coefficients were calculated at two different levels of the phenotypic data using the cor function in R with use=“pairwise.complete.obs”. First, using the BLUEs per year, correlations were calculated for all combinations of the available image-derived datasets, namely ETH24, ETH23, LfL23, LfL24 and ETH24L. Second, focusing on the field trials ETH23 and ETH24, the BLUPs resulting from Equation 4 were correlated in two different ways: within image-derived traits and between image-derived and manually measured traits. For this, two-sided *p*-values were calculated for each comparison and adjusted for multiple testing using the Benjamini–Hochberg false discovery rate at three different significance levels: *α* < 0.05, *α* < 0.01 and *α* < 0.001. Based on these values, the function hclust from stats (v.4.5.1) was used to perform hierarchical clustering to establish groups of similar traits for subsequent analysis.

#### 2.3.6. Principal component analysis

The accessions of the buckwheat collection were previously grouped into seven subclusters using NGSadmix as described in Hess et al. (2026c). In addition, the accessions were also labelled according to the morphology of their flowers (which reflects self-compatibility) and their terminal flower cluster. A principal component analysis (PCA) was performed on the centred and scaled BLUEs calculated over ETH23 and ETH24 for all 22 image-derived flowering traits using the function prcomp of the R package stats (v.4.5.1) and was used to assess differences in flowering behaviour between different subclusters and the previously determined flowering types.

#### 2.3.7. Genome-wide association studies for image-derived and manually measured flowering traits

Genome-wide association studies (GWAS) were performed using GAPIT (v.3.5.0) in R (Wang and Zhang, 2021) on image-derived and manually measured datasets from the ETH trials (2023, 2024 and combined). For each trait-dataset combination, four methods implemented in GAPIT were tested: BLINK (Huang et al., 2019), FarmCPU (X. Liu et al., 2016), mixed linear model (MLM) (Yu et al., 2006) and multi-locus mixed model (MLMM) (Segura et al., 2012). The phenotypes were year-site BLUEs, and the genotypes were supplied as an imputed allele-frequency matrix comprising 52,182 SNPs. The structure of the population was controlled using the first three principal components of the genotype matrix and relatedness was modelled using the GAPIT implementation of the genomic kinship matrix proposed by (VanRaden, 2007). For each run, marker-trait association significance was assessed using the Benjamini-Hochberg FDR at *α* < 0.05.

Two complementary approaches were used to summarize and visualize patterns of marker-trait associations from the results of 420 GWAS analyses. First, exact SNP overlaps among previously defined trait groups were visualized by generating an UpSet plot (Lex et al., 2014) from all significant marker-trait associations. Second, *p*-values of significant marker-trait associations involving the same marker across models and datasets were combined by Fisher’s combined probability test (Fisher, 1992) to produce a single combined *p*-value per position. The combined −*log*_10_(*p*) were plotted at genomic coordinates to visualize the distribution and significance of grouped marker-trait associations.

Since exact SNP overlap may miss associations originating from the same genetic signal, linkage disequilibrium (LD)-supported concordance between image-derived and manually measured GWAS results was assessed. Pairwise LD was calculated from the imputed allele-frequency matrix as dosage-based *r*^2^, defined as the squared Pearson correlation coefficient between SNP vectors across accessions (Hill & Robertson, 1968). Background-adjusted LD decay was first estimated separately for each chromosome using all SNP pairs within 2 Mb, grouped into 5 kb distance bins containing at least 100 pairs. Initial LD was estimated from pairs separated by 5 to 50 kb, whereas background LD was estimated from pairs separated by 500 kb to 2 Mb. A chromosome-specific LD window was defined as the first distance at which the LOESS-smoothed median *r*^2^ crossed the background-adjusted half-decay threshold for three consecutive bins, with a minimum window of 50 kb. These chromosome-specific LD windows (Table S2) were then used to construct LD-supported blocks by connecting SNP pairs within the corresponding window when *r*^2^ ≥ 0.20. Connected components were used as LD blocks, with their maximum span capped at the chromosome-specific LD window. Image-derived and manually measured associations were considered concordant when they involved the same SNP or when distinct SNPs occurred within the same LD block, and were classified as flowering, abscission or mixed trait-category matches.

To further investigate the associations of the image-derived phenotypes, all genes directly co-localizing with markers involved in significant marker–trait associations were retrieved. For selected associations, regional GWAS profiles were inspected across datasets and models, and local LD was visualized using the focal SNP together with 12 neighbouring SNPs on each side. Pairwise LD in these local plots was calculated from the allele-frequency matrix as described above. The predicted protein sequences of these genes were extracted from the genome assembly Tuka (Hess et al., 2026a) and used as input for EggNOG-Mapper (v.2.1.13; Cantalapiedra et al., 2021) with default parameters. This tool uses precomputed orthologous groups and phylogenies from the EggNOG (v.5.0.0) database (Huerta-Cepas et al., 2019) to transfer functional information to the queried sequences. Additionally, for all significant marker-trait associations within a gene, a linear model was fitted to model the allelic effect, where the *R*^2^ coefficient was used to describe the percentage of phenotypic variation explained.

## 3. Results

### 3.1. Image segmentation model for flower percentage estimation

During the image segmentation model training, both box and mask losses initially decreased until a steady state was reached around epoch 250 (Figure 3A). This state was maintained until the end of training. A similar pattern appeared in validation losses computed at 50-epoch intervals from saved checkpoints. The correlations between the predicted areas and the ground-truth mirrored the stability of the loss dynamics (Figure 3B), where all checkpoints showed very high correlations for both sets (*r* > 0.97). The unprocessed predicted masks from the training set showed a higher correlation with the ground truth compared to the validation set. After merging overlapping masks, all the correlations increased to *r* > 0.99, indicating that the raw output occasionally predicted the same mask multiple times. The best-performing checkpoint based on correlation (after merging the masks) occurred at epoch 550 and is shown in detail in Figure 3C. The scatterplot for this checkpoint shows a slight underprediction of Flower %, visible as a small shift of the points from the 1:1 line. This bias is negligible for downstream analyses because it is systematic across images processed by the same model and does not affect the overall distribution. Images without annotated flowers showed an average predicted flower area of 0.5%, with some up to 3% (Figure 3D). Based on this, a percentage of 2% was set as the detection limit for our trained image segmentation model.

**Figure 3.**
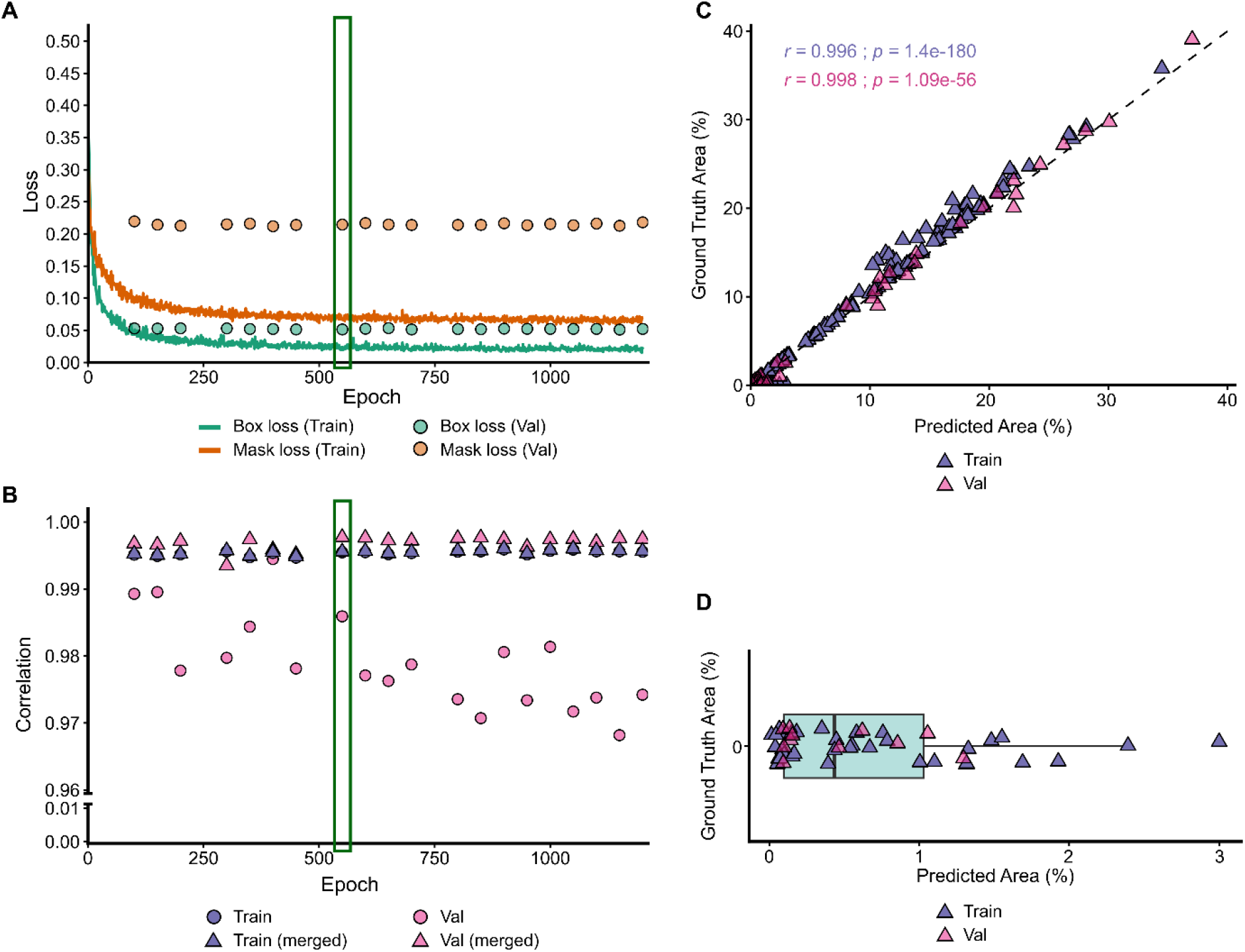
Training and validation metrics for the buckwheat flower image segmentation model. (A) Evolution of box (orange) and mask (green) losses over training for the training set (lines) and the validation checkpoints (points). (B) Correlations between predicted and ground truth area across training checkpoints for the train (purple) and validation (pink) sets for the raw mask predictions (circles) and the merged overlapping masks (triangles), note that the y-axis is truncated for a better visualization. Green boxes in A and B depict the checkpoint at which the best metrics were attained. (C) Scatterplot of Predicted Merged Area (%) and Ground Truth Area (%) for the selected best model depicting both training (purple triangles) and validation (pink triangles) sets. (D) Box plot of predicted area (%) for images with a ground truth area (%) equal to zero.

### 3.2. Modelling of buckwheat flowering dynamics

Compared to the other experiments, the ETH23 and ETH24 trials were characterized by a higher data density, with a median of 22 and 31 images per plot and time-series durations of 95 and 121 days on average for 2023 and 2024, respectively (Table 1). All the datasets showed an overall satisfactory goodness-of-fit, with median *R*^2^ values above 0.85 for both the flowering and abscission phases across all experiments (Figure 4A). The three-parameter logistic function describing the flowering phase achieved higher goodness-of-fit than the linear function used for the abscission phase, with median *R*^2^ values ranging from 0.98 to 0.99 and from 0.85 to 0.91, respectively. After filtering for fits with *R*^2^ > 0.5 three plots were excluded for the flowering phase and 35 plots for the abscission phase, and their corresponding parameters were discarded (Table 1). Notably, the ETH23 trial showed the highest number of discarded abscission-phase models, with 35 removed out of 594. An example of model fits for a single accession (FE 268) across all datasets is shown in Figure 4B, illustrating some of the contrasting metrics presented in Table 1, such as plots per accession, number of time points and mean duration.

**Figure 4.**
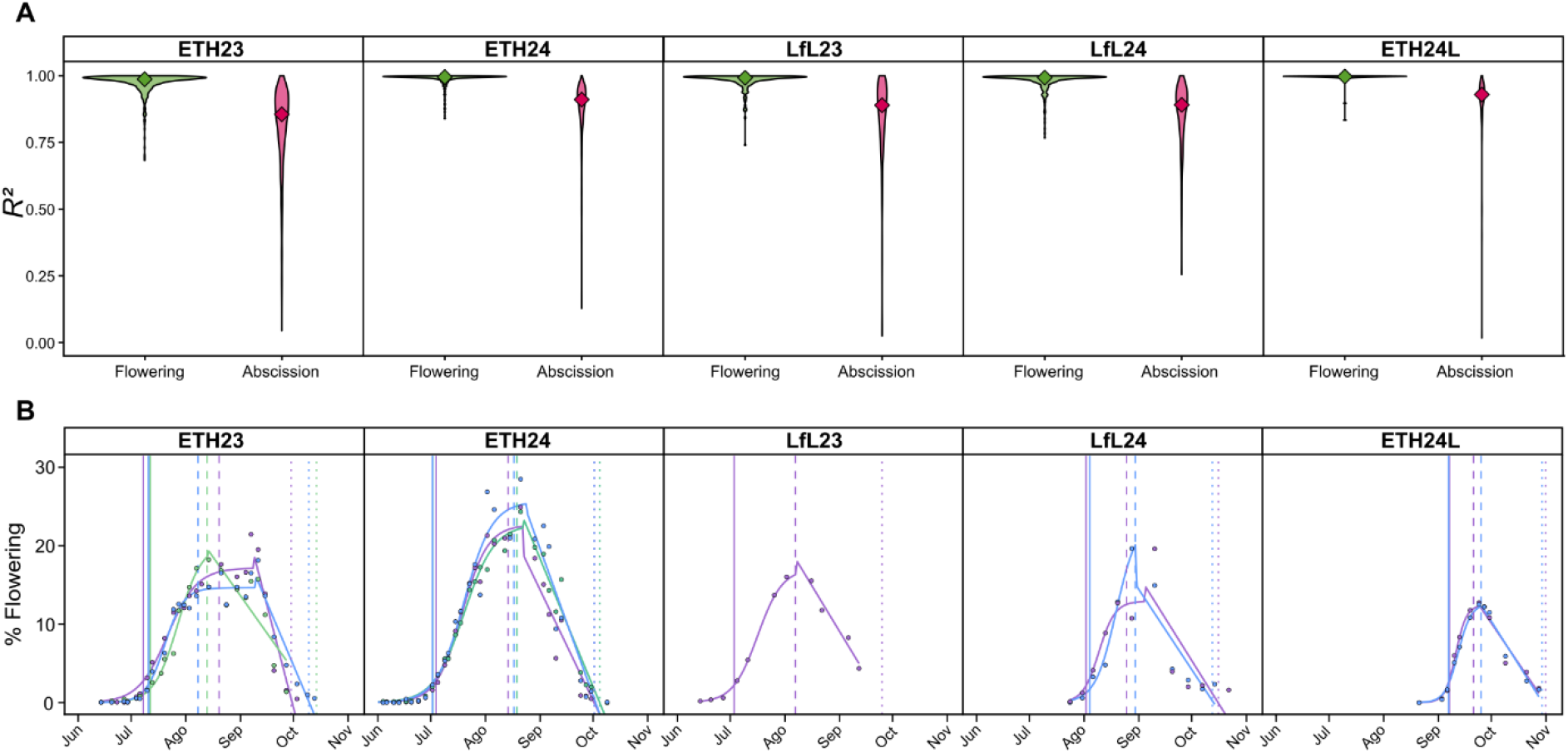
Flowering dynamics modelling. (A) Violin plots for the goodness-of-fit of both flowering phases at the field plot level in the basis of *R*^2^ for the five used datasets: ETH23, ETH24, LfL23, LfL24 and ETH24L. Diamonds represent the median of the distribution. Green and magenta represent the flowering and abscission phases, respectively. (B) Example of model fits for the accession FE268 across all datasets. Different colours represent different plots for the same accession. The solid lines represent the different Start Flowering DAS, the dashed lines the End Flowering DAS and the dotted lines the Projected End of Abscission DAS for each plot

### 3.3. Variation in flowering traits across sowing time, accessions and environments

Buckwheat grown as a second crop at ETH (ETH24L) reached all flowering phases earlier (days after sowing) than buckwheat grown as a main crop (combined ETH23 and ETH24; Figure 5A1). Flowering intensity and Flowering AUC were also lower in the second-crop trial (Figures 5A2–A3), and all differences were statistically significant (Wilcoxon-test: *p<*0.001). Moreover, the variation between accessions of buckwheat as a second crop was lower for time point traits (Figure 5A1) and Flowering AUC (Figure 5A3) while this pattern was not observed for flowering intensity traits. For example, on average buckwheat sown as the main crop reached Projected End of Abscission DAS after 124 days, and the trait showed a standard deviation of 15.86 days, while the buckwheat sown after wheat reached the same development stage already after 79 days, and the standard deviation among accessions was only 7.03 days. These patterns as well as the significant differences between buckwheat grown as a main crop and as a second crop remained persistent when only overlapping accessions between the trials were considered.

The buckwheat collection analysed in this study is composed of several subclusters (Hess et al., 2026b) and contains a total of nine self-compatible and eleven accessions with determinate growth habit. These different subclusters showed distinct flowering dynamics as shown in the PCA (based on combined data from ETH23 and ETH24) in Figure 5B. Accessions from subclusters 1 (mainly originating from East Asia) and 7 (diverse origin) were associated with late Start Flowering DAS and Projected End of Abscission DAS while subcluster 3, which includes accessions from the Himalayan region, showed moderately late flowering and slow Abscission Rate (higher, i.e., less negative values for abscission slope indicate a slower decline during abscission phase). Subclusters 2 and 4, representing Central and Eastern accessions, respectively, showed early and short generative development. Determinate and self-compatible accessions (from subcluster 5) both showed low flowering intensity and low Flowering AUC.

**Figure 5.**
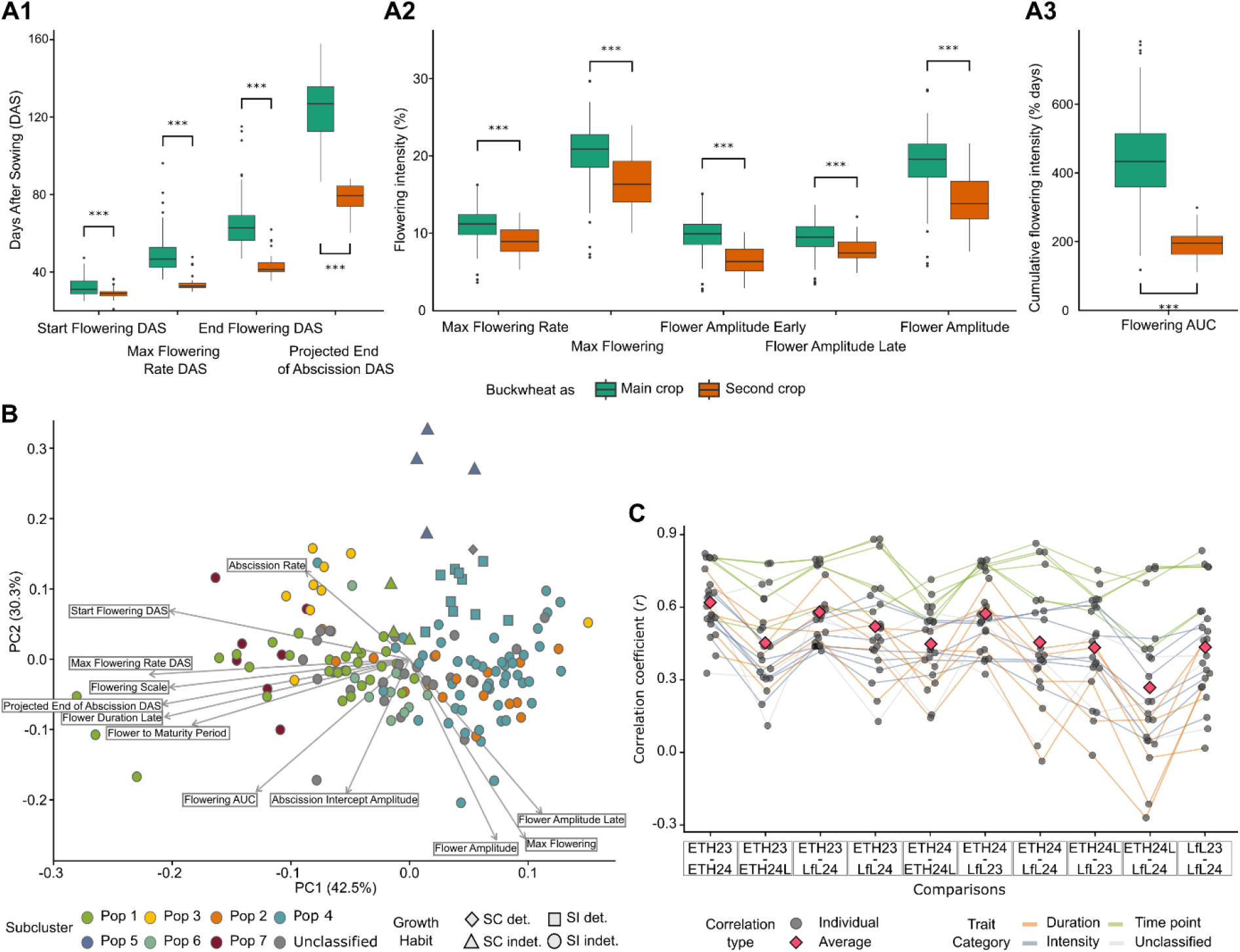
Effects of sowing time, accessions and environments on the flowering dynamics of buckwheat. (A1-3) Boxplots showing the range of flowering traits within the ETH Zurich/BLW buckwheat collection sown as main (green; *n* = 167) or second crop (orange; *n* = 51). The three plots show (1) traits related to time points in flower development, (2) flowering intensity and (3) Flowering AUC. Stars indicate significant (*p* < 0.01, Wilcoxon-test) differences between parameters compared. (B) Principal component analysis revealing differentiation of subclusters of buckwheat in terms of flowering traits. For readability, only a selection of traits is shown in the figure, while the principal components are based on all 22 image-derived flowering traits. Colours represent subclusters, and shapes growth habit. (C) Pairwise comparisons in terms of Pearson’s correlation coefficient (*r*) between environments of the 22 image-derived flowering traits. The different line colours indicate traits related to duration (orange), intensity (blue) and timing (green) of flowering phases, and the pink diamonds indicate the mean over all 22 traits per year-site comparison.

Comparison of pairwise correlation coefficients for flowering traits observed in different environments shows that some traits are highly correlated in all comparisons, while others have consistently low correlations (Figure 5C). Traits related to time points such as Mid flowering DAS, Max Flowering Rate DAS, Start Flowering DAS or Projected End of Abscission DAS showed the highest average correlation over all year-site comparisons (0.64 < *r* < 0.77). Correlations between year-sites were lower (*r* < 0.6) for traits describing durations of flowering phases or flowering amplitudes. Low correlations (*r* < 0.4) were also recorded for Abscission Rate and Flowering AUC. Comparing the average correlation coefficients between the year-site comparisons revealed that correlations between the trial with buckwheat as a second crop with trials where buckwheat was planted as a main crop were weaker (*r̅* < 0.5) compared to correlations between main crop trials (*r̅* > 0.5, except for the comparison LfL23 and LfL24). The highest average correlations coefficients were found for the comparison ETH23 and ETH24 with *r̅* equal to 0.62. However, even for comparisons of year-sites with different sowing times, correlations for time point traits remained high with values above 0.6 (*r*). The same pattern as for the correlations between year-sites was observed in broad-sense heritabilities, i.e., heritability was higher for traits related to time points compared to traits describing the durations of phenological phases or flowering amplitudes (Tables S3-S5).

### 3.4. Comparison of image-derived and manually measured flowering traits

Given the availability of BBCH traits scored manually for the ETH23 and ETH24 trials (Table S6), together with the high temporal resolution used to extract image-derived flowering phenotypes, these datasets were used for subsequent comparisons between both approaches. The pairwise Pearson correlations of the BLUPs over years between image-derived traits presented high values of *r* within themselves and broadly fell into three major groups (Figure 6A). The phenotypes describing the Flower % (flowering *y*-axis) highly correlated among themselves, with values ranging from 0.77 to 1 (mean of 0.92), and constituted a clear group named hereafter Image-derived Flower Value. The group describing time points as well as periods during the flowering phase (flowering *x*-axis) correlated with high to modest *r* values with an average (*r̅*) of 0.69 in a group named Image-derived Flower Period. The last group of phenotypes, Image-derived Abscission, was constituted by most of the parameters extracted from the abscission modelling phase that also presented high *r* values with *r̅* = 0.78. It is important to note that the traits Abscission Intercept Amplitude and Flowering AUC could have been assigned to other groups given their meaning and *r* values but were assigned to the Image-derived Flower Period group for their overall higher *r* with the rest of the traits in this group.

**Figure 6.**
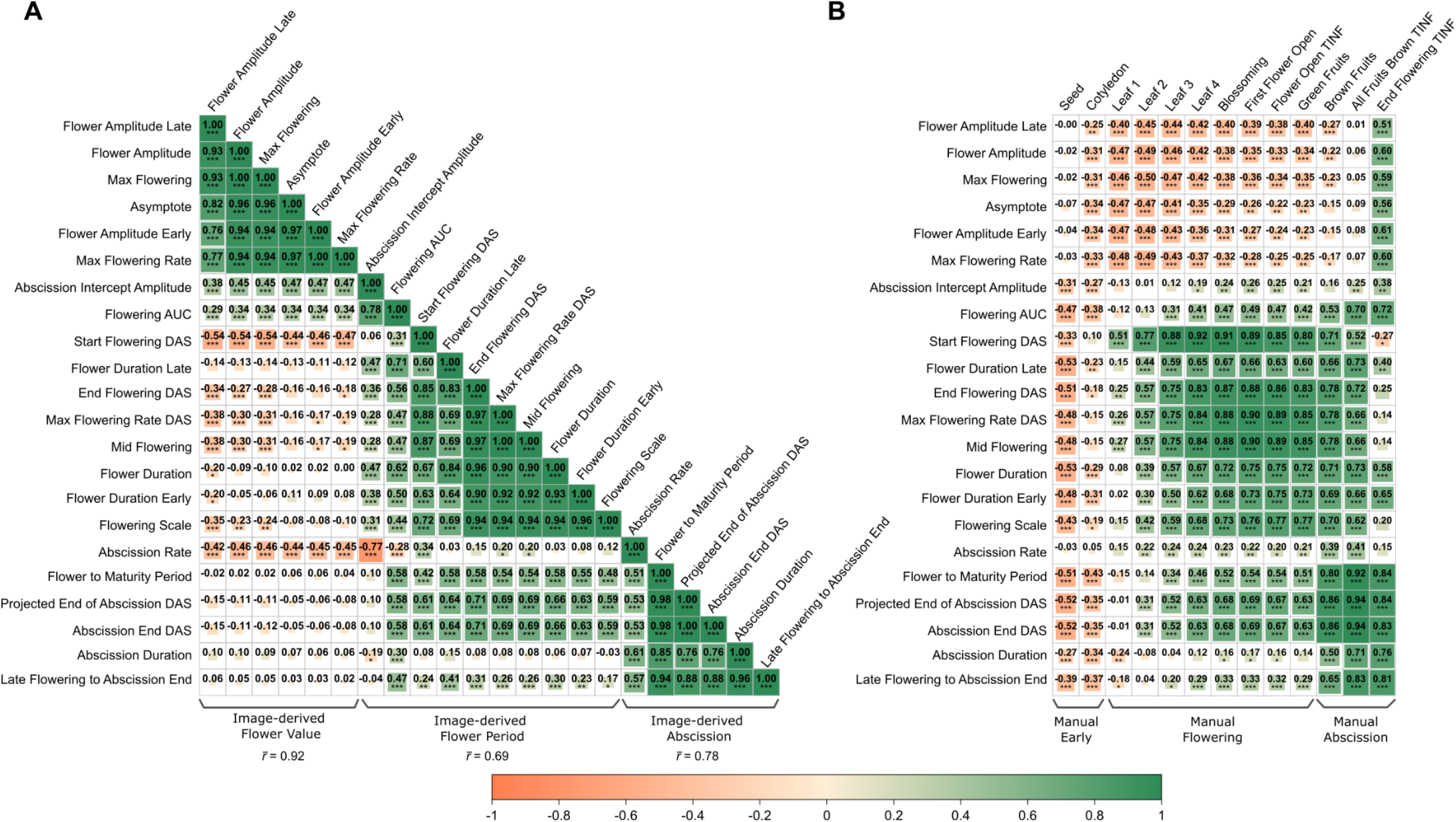
Pairwise Pearson correlations (*r*) between BLUPs across years of ETH main experiments (A) within image-derived traits (B) and between image-derived and measured traits. Positive correlations are depicted in green and negative correlations in orange, with a saturation and a square size proportional to their absolute correlation (|*r*|). Asterisks indicate the significance from two-sided correlation tests at different significance levels: ^∗^*p* < 0.05, ^∗∗^ *p* < 0.01 and ^∗∗∗^ *p* < 0.001. In (A) *r̅* indicates the mean pairwise correlations within each group excluding self-correlations.

When comparing image-derived phenotypes with temporally ordered BBCH stages, additional patterns were revealed (Figure 6B). The first image-derived group (Image-derived Flower Value) showed a consistent correlation pattern with the manually measured traits, with a weak but significant negative correlation with values ranging from -0.17 to -0.50 with the stages of vegetative development, and a weak but significant correlation ranging from 0.51 to 0.6 with End Flowering TINF, which describes the end of flowering in the terminal inflorescence. All correlations between manually measured traits and image-derived traits in this group were below |*r*| = 0.6.

The second image-derived group (Image-derived Flower Period) also presented a consistent correlation pattern with the manually measured BBCH stages during flowering development (Manual Flowering). Timepoint-related phenotypes, such as Start Flowering DAS, Max Flowering Rate DAS, and Mid Flowering showed the highest correlations with up to *r* = 0.92, whereas period-related phenotypes presented values around 0.75. Finally, the third group of image-derived phenotypes related to abscission (Image-derived Abscission) also showed a consistent pattern of correlations with the manually measured BBCH stages. These traits showed negative correlations with early BBCH stages starting at *r* = -0.52, which increased along the subsequent stages with some values up to 0.69. Highest correlations were found with the last measured BBCH stages (Manual Abscission) with values ranging from 0.50 to 0.94. It has to be noted that Abscission Rate, although following the same pattern, presented an overall weaker correlation with all the BBCH stages compared to the other phenotypes from the Image-derived Abscission group.

### 3.5. Comparison of genome-wide association studies between image-derived and manually measured traits

Image-derived phenotyping methods generated a higher number of associations (*n* = 239) compared to manual scoring methods (*n* = 170) across the ETH23, ETH24, and combined datasets when comparing Flower Period and Abscission traits (Figure 7A). Most of these associations originated from the Image-derived Flower Period group (*n* = 197), which is the one constituted by the highest number of traits (*n_t_* = 10). Within these groups, only three significant associations were shared between Image-derived Flower Period and Image-derived Abscission. For BBCH-based phenotyping scores, a similar trend was observed, with Manual Flowering as the group with the most associations (*n* = 118), nine of which were shared with the group Manual Abscission. Generally, 26 associations involved the same markers between the phenotyping methods, with 16 of these markers present in both Image-derived Flower Period and Manual Flowering. A general overview of the 463 significant marker-trait associations for both phenotyping methods in their chromosomal context is displayed in Figure 7B. The 26 direct common associations were found in six of the eight chromosomes (Chr). Chr3, Chr7 and Chr8 harbour the highest number of associations (*n* = 21). Accounting for LD between nearby markers 21 additional coincidences between image-derived and manually measured GWAS associations were identified. These coincidences mostly clustered around loci already detected through exact SNP overlap, rather than defining new signals. Only four pointed to additional shared regions without a nearby direct overlap, located on Chr1, Chr2 and Chr6. Additional regions without direct or LD-supported coincidence but with close associations between methods can also be observed. Some interesting regions fulfilling these criteria are the first megabase pairs from Chr6, the last from Chr1, or around the position 50Mb on Chr4. Figures S1–S2 report the complete set of marker-trait associations for each trait across datasets and GWAS models.

**Figure 7.**
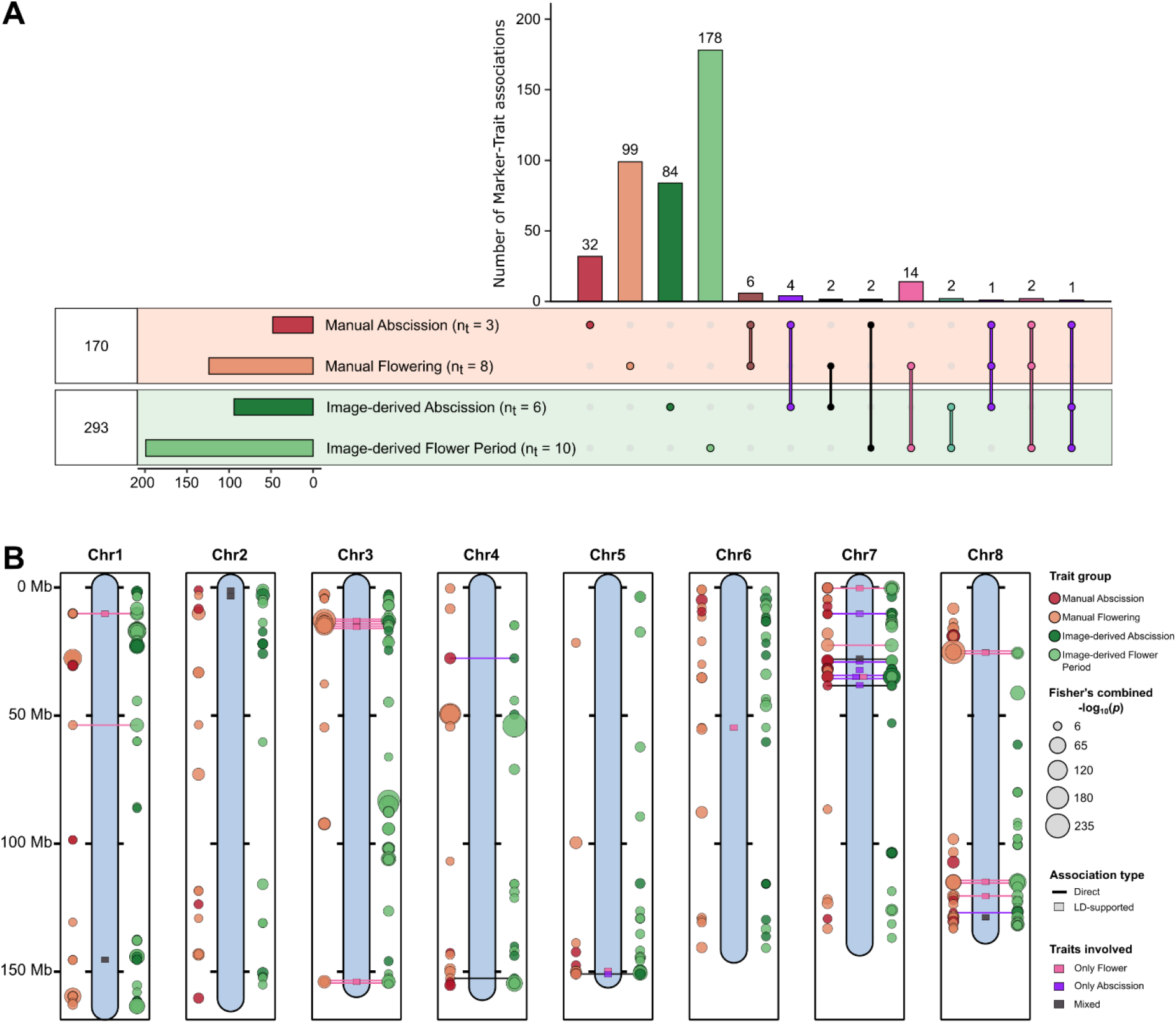
Concordance of marker–trait associations between image-derived and measured flowering traits. (A) UpSet plot showing unique and shared significant markers among four trait groups: Manual Abscission, Manual Flowering, Image-derived Abscission and Image-derived Flower Period. The number of traits included in each group is indicated by *n_t_*, the horizontal bars show the number of unique markers in each group, and the numbers at the left indicate the total number of unique markers identified for each phenotyping method. Intersections involving both phenotyping methods are classified according to whether the shared association concerns flowering traits (pink), abscission traits (purple) or both trait categories (black). (B) Genomic distribution of significant marker–trait associations and concordant signals across the eight chromosomes. Manually measured trait associations are shown to the left of each chromosome and image-derived trait associations to the right. Point colour denotes the trait group, while point size is proportional to Fisher’s combined −*log*_10_(*p*) for associations located at the same genomic position. Direct associations denoted by horizontal bars connect identical SNP positions detected by both phenotyping methods. LD-supported associations denoted by squares indicate concordance between distinct SNP positions located within the same LD block. Pink, purple and black denote associations involving only flowering traits, only abscission traits and both trait categories, respectively.

### 3.6. Putative candidate genes governing flowering dynamics in buckwheat

The genetic context of the Image-derived phenotype associations was examined in detail using the uniquely identified 343 marker-trait associations. Of these, 265 were found to co-localize with 171 annotated genes, 93 of which contained markers located within the putative coding sequence (CDS; Table 2). Four gene-overlapping marker-trait associations were selected for detailed inspection using regional GWAS profiles, local LD heatmaps and marker-effect plots (Figure 8). Across the four loci, the regional GWAS profiles showed localized association peaks at the focal markers. The local LD heatmaps showed neighbouring markers with variable pairwise *r*^2^, but did not show continuous high-LD blocks spanning the full plotted intervals. The marker “chr1_163426249_G/A” affecting Mid Flowering, co-localizes with the gene “evm.TU.chr1.4316”, annotated as an “Auxin response factor (ARF)” (Figure 8A). This marker had a strong and significant effect across all three datasets, with *R*^2^ values of 0.374, 0.351 and 0.397, indicating that a substantial proportion of the trait variance was explained by this single marker. The marker “chr8_79940130_C/T” affecting Abscission Rate, co-localizes with the gene “evm.TU.chr8.992” annotated as “Ethylene-insensitive protein” (Figure 8B). The marker “chr6_16913820_G/C” affecting Flowering Amplitude, is located within the gene “evm.TU.chr6.1780” annotated as a “Nuclear chaperone” (Figure 8C). Finally, the marker “chr7_633641_G/A” affecting Start Flowering DAS, co-localizes with the gene “evm.TU.chr7.98” annotated as a “Diphthamide biosynthesis protein” (Figure 8D). The latter three markers showed weaker effects in ETH23, with *R*^2^ values ranging from 0.048 to 0.199, when compared to the ETH24 and combined datasets, with *R*^2^ values between 0.213 and 0.322. Further details showing all marker-trait associations that fell within annotated genes across the three datasets, are provided in Table S7.

**Figure 8.**
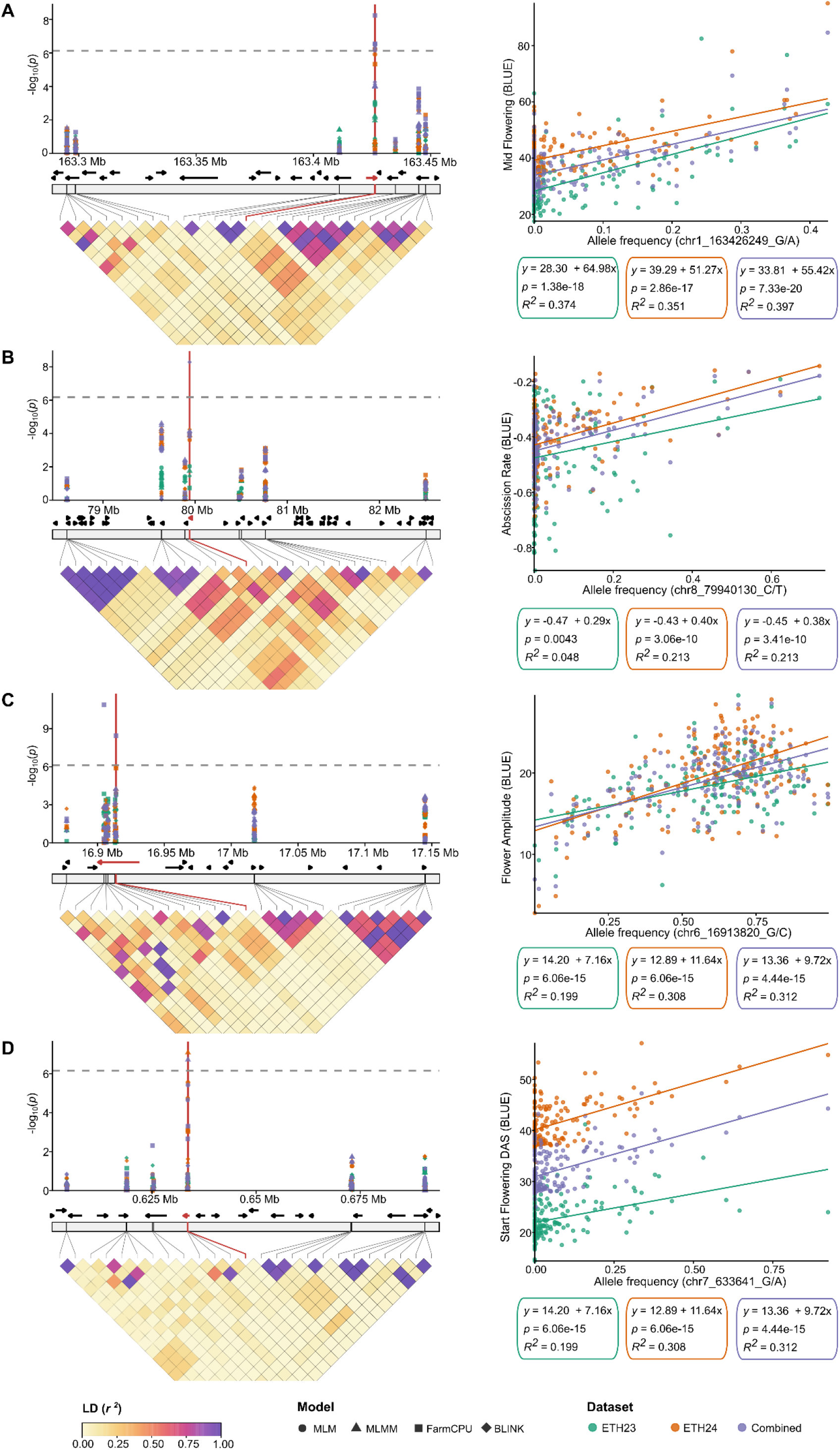
Regional association patterns, linkage disequilibrium and allelic effects for four selected marker–trait associations. (A) chr1_163426249_G/A associated with Mid Flowering. (B) chr8_79940130_C/T associated with Abscission Rate. (C) chr6_16913820_G/C associated with Flower Amplitude. (D) chr7_633641_G/A associated with Start Flowering DAS. For each locus, the left panel shows a regional association plot based on physical genomic position for ETH23, ETH24 and the combined dataset, using four GWAS models: MLM, MLMM, FarmCPU and BLINK. The dashed horizontal line indicates the significance threshold, and the red vertical line marks the focal SNP. For the LD analysis, the focal SNP and its 12 nearest neighbouring SNPs on either side are shown. Gene annotations are displayed below the regional association plot, with black arrows indicating transcriptional orientation and the gene containing the focal SNP highlighted in red. The selected SNPs are connected to the triangular LD heatmap, which displays pairwise linkage disequilibrium as *r*^2^. The right panel shows the relationship between allele frequency at the focal SNP and trait BLUEs for ETH23, ETH24 and the combined dataset. Coloured lines represent dataset-specific linear fits, and the boxes report the fitted equation, the significance of the slope (*p*) and the proportion of variance explained (*R*^2^).

**Table 2.** Overview of marker-trait associations and their co-localization with annotated genes. The total number of marker-trait associations, number of associations within a gene, unique number of genes associated to traits as well as how many of them present an association in the coding sequence (CDS).

| **Group** | **Markers (n)** | **Markers in gene (n)** | **Unique genes (n)** | **Genes with CDS hit (n)** |
|---|---|---|---|---|
| Image-derived Flower Period | 199 | 150 | 98 | 63 |
| Image-derived Flower Value | 53 | 44 | 34 | 13 |
| Image-derived Abscission | 94 | 73 | 43 | 21 |
| Image-derived all | 343 | 265 | 171 | 95 |

## 4. Discussion

Our image-based segmentation approach yielded accurate predictions of Flower %, with a strong correlation between predicted and manually measured values (*r* > 0.99). The robustness of the image-derived flowering traits describing buckwheat flowering patterns was further supported by high repeatability and heritability for most traits. In addition, the image-derived traits accurately predict all manually measured BBCH stages related to flowering (*r* > 0.9) and seed formation (*r* > 0.85), demonstrating their biological and agronomic relevance. Most importantly, our phenotyping approach encompassed different sowing times (main and second crop), environmental conditions (sites and years), imaging devices (LITERAL®, smartphone, high resolution digital camera) and field staff responsible for image acquisition. This demonstrates that the workflow described here (image-acquisition, image-segmentation, quantification of flowering dynamics) represents a robust and broadly applicable phenotyping protocol. This robustness is especially relevant as multi-environment trials are common in variety testing (Isik et al., 2017) and typically consist of trials spread over distant locations that do not have access to the same equipment and field personnel. Therefore, image-based phenotyping enables the quantification of flowering traits in an objective manner, improving the comparability of data between trials conducted under various conditions.

The three-parameter logistic model used to describe the flowering phase, and the linear model applied to the abscission phase showed good model fits, confirming their suitability for capturing major developmental trends. In contrast, poorer model fits were more frequently observed for the abscission phase for the ETH23 trial. This was associated with a second flowering peak in some plots towards the end of August 2023, triggered by a specific weather pattern characterized by a prolonged period of hot and dry weather followed by five days of heavy rain (Supplementary Figure 3). No comparable environmental conditions were observed in summer 2024 (Supplementary Figure 4) and no second flowering peak occurred. This indicates a limitation of the modelling approach rather than data quality *per se*, as the assumption of a single linear trajectory does not hold true under all environmental conditions. Consequently, future studies based on larger datasets collected across a wider range of environmentsmay benefit from more flexible modelling approaches to better capture complex flowering dynamics.

### 4.1. Image-based, high throughput phenotyping of buckwheat flowering patterns

Vegetative development stages assessed using the BBCH scale were most strongly correlated with Start Flowering DAS, the image-derived flowering trait that is temporally closest to vegetative stage. This correlation was strongest for later vegetative phases (e.g., formation of the fourth true leaf) and weaker for earlier stages such as cotyledon formation. This pattern is expected, as the image-based analysis focused on floral development rather than vegetative growth. Thus, our imaging approach complements efforts to optimise BBCH scales for buckwheat (Arduini et al., 2016; Grimes et al., 2025) by providing an automated workflow for assessing flowering dynamics in large scale field trials and variety testing. In particular, it delivers high-quality phenotypic data for the period from the onset of flowering to the early seed set, which has been described as the most critical period for seed formation (Guglielmini et al., 2019).

In addition to timing of flowering, the image-based trait detection provides quantitative information on flowering intensity, which is not captured by the BBCH scale and, to our knowledge, not previously assessed under field conditions. Traits related to flowering intensity have been shown to be relevant for plant breeding in other crops (Zhang et al., 2020, Oppliger et al., 2025), and combining Flowering AUC with the yield data could serve as a proxy for expected yield. Most promising would be the development of a metric to assess seed set efficiency (i.e., the proportion of flowers that produce seeds) under field conditions. Although seed set efficiency is considered a major limitation for buckwheat productivity (Campbell, 1997), its assessment has been constrained by the difficultly of quantifying flower numbers in the field. This limitation could be addressed by adapting the methods described here. Beyond seed formation, flowering intensity is also relevant if the plant is used in ecological flower strips (Strahm et al., 2019) or as a melliferous crop (Halbrecq et al., 2005), further illustrating the broader relevance of flowering intensity traits.

### 4.2. Variation of phenology in the buckwheat collection

Very little data on the flowering behaviour of different buckwheat accessions is available. However, Campbell (1997) reported data from variety testing in India (number of accessions = 408), Nepal (*n* = 309) and China (*n* = 964), including observations on the start of flowering, full flowering and maturity. Comparison between these data and our results must be made with caution due to differences in the definitions of development stages and the lack of metadata about the trials, such as environmental conditions at the trial sites and information on sowing time. Strahm et al. (2019) reported the start and full flowering of 14 accessions tested in Switzerland under similar climatic conditions to those of our trials. The mean values reported for start of flowering, full flowering and maturity are generally comparable to those found here. However, our data cover a wider range of values, which likely reflects the greater genetic diversity of the accessions investigated.

The diversity of flowering behaviours contained in our buckwheat collection represents a resource to be harnessed to breed new buckwheat varieties tailored to fulfil specific roles in the crop rotation. In the literature, buckwheat has been suggested as a main or second crop for grain production (Radics and Mikóházi, 2010), a fodder crop (Amelchanka et al., 2010), a green manure (Taranenko et al., 2016) or a melliferous flower crop (Halbrecq et al., 2005). Each of these roles requires distinct flowering characteristics. For example, varieties intended as a main crop for grain production should be bred to maximize seed set, whereas varieties developed for green manure may benefit from limited seed set, to reduce volunteer plants in subsequent crops (McKenzie-Gopsill, 2025). Varieties used as a second crop must exhibit rapid development and reach maturity before the first frost, while melliferous varieties would be selected for sustained flower production (i.e., maximize Flowering AUC or Flower Amplitude). Across all applications, a rapid and homogeneous abscission phase (i.e., steep Abscission Rate and short Abscission Duration) would make harvesting and processing easier (Funatsuki et al., 2000; Arduini et al., 2016).

However, our results also show that sowing time (i.e., different day length during development) has a large effect on buckwheat flowering behaviour as previously reported (Hagiwara et al., 1998; Michiyama et al., 2005; Arduini et al., 2016). Therefore, it is crucial to assess the response of different accessions to changing day length and to perform variety testing and selection under environmental conditions similar to their intended production location and system (Ewing et al., 2019).

### 4.3. From pictures to genes

In total, 343 unique markers were associated with one or more of the 22 image-derived flowering phenology traits, reflecting the complexity of such traits as reported for other crops (C. Liu et al., 2021; Shi et al., 2022; Y. Li et al., 2023). Of these associations, 171 occurred within annotated genes, offering insights into the possible molecular mechanisms underlying the variation detected. The regional profiles generated for selected gene-overlapping markers showed localized association peaks and no broad continuous high-LD blocks across the neighbouring markers. This pattern is consistent with the rapid LD decay expected in outcrossing species and with previous reports of low LD in allogamous common buckwheat populations (Flint-Garcia et al., 2003; Yabe et al., 2018; Matsui and Yasui, 2020). LD was calculated from genome-wide allele-frequency fingerprints, meaning that the local *r*^2^ heatmaps describe correlations among accession-level allele-frequency profiles rather than individual haplotypes (Nay et al., 2020). These gene-overlapping associations, together with their functional annotations and regional GWAS profiles, highlight candidates that could be considered for different breeding objectives.

For example, a marker associated with Flowering Start DAS lies within a DPH1 homolog and is annotated as introducing a premature stop codon. *Arabidopsis dph1* mutants show delayed development and reduced cellular proliferation (H. Zhang et al., 2022). Similarly, the marker associated with Mid Flowering mapped within an auxin response factor 2 (ARF2) homolog. In *Arabidopsis*, *arf2* mutants show altered flowering time (Ellis et al., 2005; Okushima et al., 2005), supporting this locus as a plausible regulator of flowering progression. Together, these loci represent promising candidates for marker-assisted selection or introgression aimed at adjusting flowering onset or synchrony.

In contrast, the marker associated with Flowering Amplitude is directly relevant for breeding melliferous ideotypes, as it directly affects the flowering intensity during the growing season. This marker lies within a MDN1 homolog, which affects flowering time and auxin homeostasis in weak *Arabidopsis* mutants (K. Li et al., 2021, 2022), making it a candidate for flower development and therefore nectar availability. Finally, the marker associated with Abscission Rate is relevant for ideotypes targeting the persistence versus rapid decline of floral display after peak flowering. This locus lies within a homolog of an ethylene signalling gene (EIN2) in *Arabidopsis*. This gene is associated with floral organ abscission and senescence phenotypes in *Arabidopsis* and tobacco (Patterson and Bleecker, 2004; Chakrabarti and Bharti, 2023), suggesting a role in flower retention dynamics. Depending on the breeding target, this association could be explored to either prolong flowering (e.g. melliferous) or to promote a faster abscission (e.g., synchronize maturity, improve management timing). While these highlighted examples illustrate specific markers associated with flowering time, flower intensity and abscission, many additional associations exist within each of these categories (Table S7). These results provide a catalogue of candidate loci and genes that can now be prioritized for functional validation and converted into molecular tools to tailor common buckwheat varieties for target flowering dynamics and end-use ideotypes.

## 5. Conclusion

This work illustrates how image-based phenotyping coupled with statistical modelling can offer a robust and scalable platform to detect flowering traits in buckwheat across different environments, divergent varieties and phenotyping setups, including low-cost imaging devices such as smartphones and digital cameras. The resulting traits have high heritability and repeatability and correlate highly with manually measured BBCH flowering stages validating their reliability. Moreover, the ability to quantify flowering intensity adds a complementary layer of information not available in BBCH data. Thus, our phenotyping approach has the potential to accelerate buckwheat breeding and improve its flowering behaviour, overcoming a major hurdle to the crop’s adoption by farmers.

## Data availability

Code and data will be made available upon reasonable request.

## Authors contributions

F.H., L.K., S.Y. conceived this study and B.S. acquired funding. F.H. designed the field trials and managed the trials at ETH, took images (ETH only), contributed to statistical analysis and wrote the manuscript. C.Q.T, designed and implemented the image analysis pipeline, compiled the training dataset, contributed to statistical analysis, performed GWAS and wrote the manuscript. F.T. provided the scripts for the modelling of flowering dynamics. G.R. managed the trials at LfL and took images. S.Y., F.T., L.K. and B.S. edited the manuscript.

## Acknowledgements

We would like to sincerely thank Heidi Heuberger for providing administrative and scientific support to the project as the responsible coordinator of the trial at LfL. Additionally, we would like to thank Simon Corrado for his support with the establishment and management of the field trials at ETH as well as our interns and student assistants Shuyo Nodasaka, Benjamim Fabri and Manuel Cavigelli for their work in maintaining field trials and taking images.

