## Supplementary Figures for "From smartphone to gene: Automated image-based phenotyping captures flowering diversity of common buckwheat"

### Supplementary Tables

Supplementary tables are provided in the file “SupplementaryTables1-7.xlsx”

### Supplementary Figures

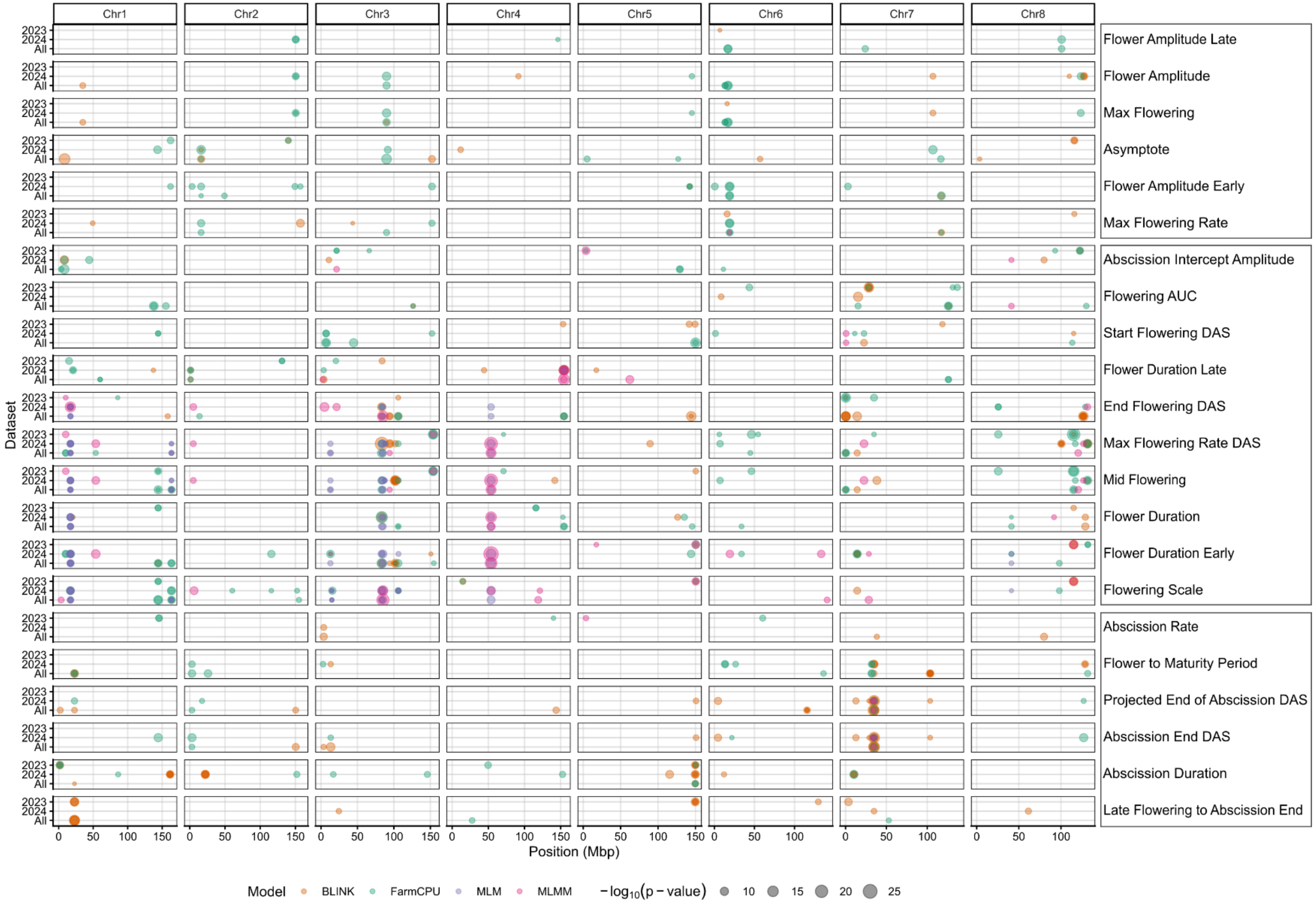


**Supplementary Figure 1.** Overview of significant marker-trait associations for the image-derived flowering phenotypes in the datasets ETH23, ETH24 and combined. Each row is a trait, each column a chromosome with the axis in mega base-pairs (Mbp). Significant markers with significant associations are colour-coded according to the method through which they were identified: BLINK (Orange), FarmCPU (Green), MLM (Purple) and MLMM (Pink) and with a size proportional to the associated $-log_{10}(p)$.

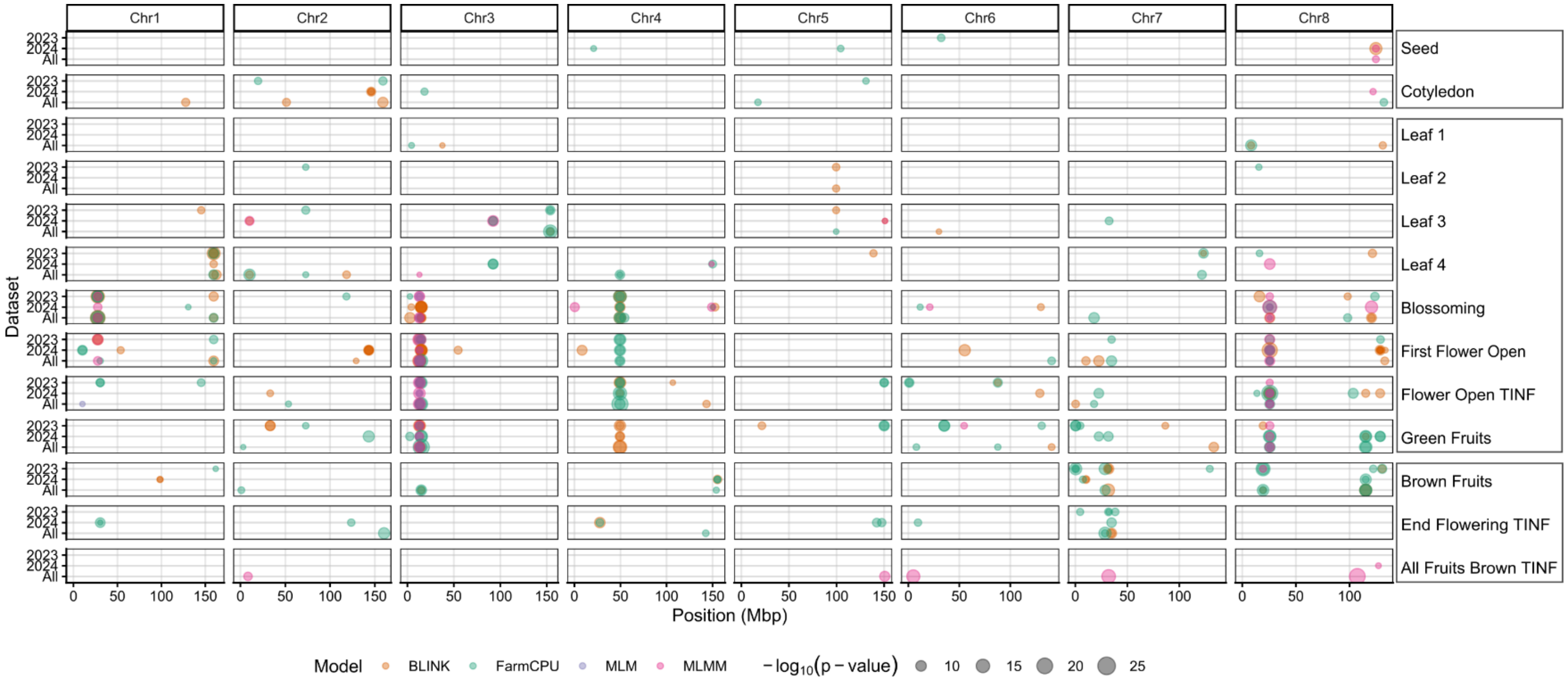


**Supplementary Figure 2** Overview of significant marker-trait associations for the manually measured flowering phenotypes in the datasets ETH 2023, ETH 2024 and combined. Each row is a trait, each column a chromosome with the axis in mega base-pairs (Mbp). Significant marker traits are colour-coded according to the method through which they were identified: BLINK (Orange), FarmCPU (Green), MLM (Purple) and MLMM (Pink) and with a size proportional to the associated $-log_{10}(p)$.

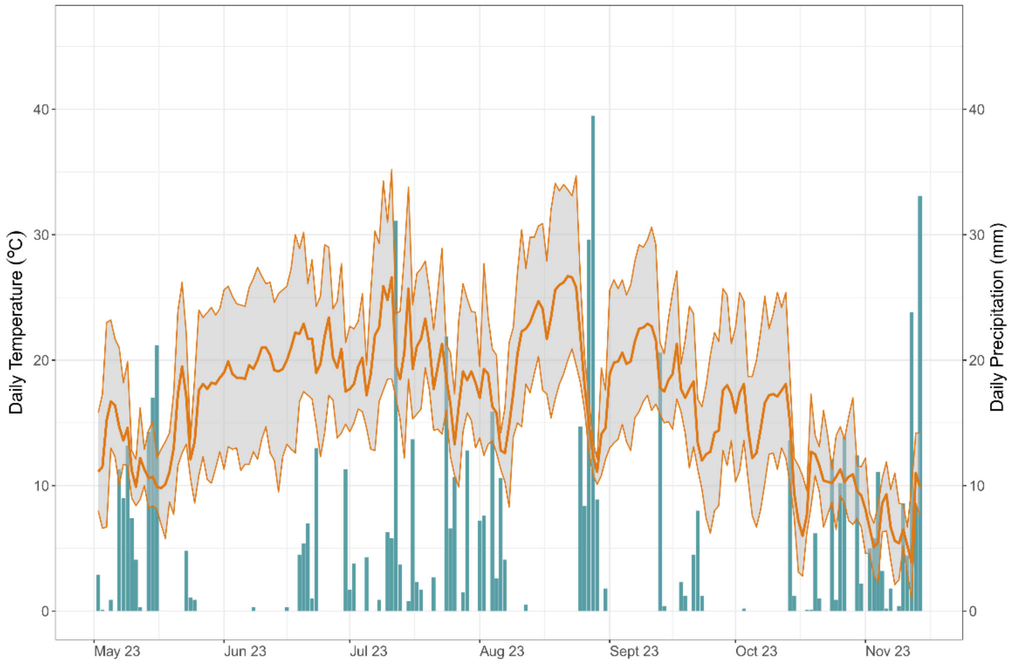


**Supplementary Figure 3.** Weather data including temperature and precipitation during the buckwheat growing season 2023 in Eschikon (ETH23). Orange lines represent daily temperatures with the bold line indicate daily mean values while thin lines show minimum and maximum temperatures. The blue bars indicate daily precipitation.

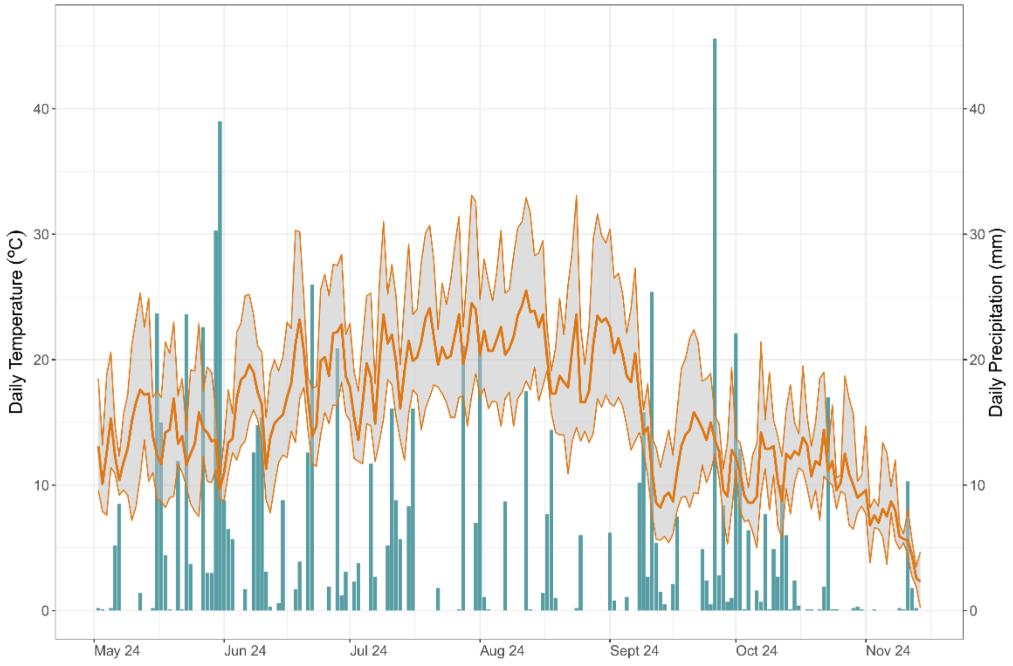


**Supplementary Figure 4.** Weather data including temperature and precipitation during the buckwheat growing season 2024 in Eschikon (ETH24). Orange lines represent daily temperatures with the bold line indicate daily mean values while thin lines show minimum and maximum temperatures. The blue bars indicate daily precipitation.
